# Phenotypic plasticity is aligned with phenological adaptation on both micro- and macroevolutionary timescales

**DOI:** 10.1101/2021.01.26.428241

**Authors:** Stephen P. De Lisle, Maarit I. Mäenpää, Erik I. Svensson

## Abstract

In seasonally-variable environments, phenotypic plasticity in phenology may be critical for adaptation to fluctuating environmental (temperature) conditions. Using an 18-generation longitudinal dataset from natural insect (damselfly) populations, we show that phenology has strongly advanced. Individual fitness data suggest this is likely an adaptive response towards a thermally-dependent fitness optimum. A laboratory experiment revealed that developmental plasticity qualitatively matches environmental-dependence of selection, partially explaining observed phenological advance. Expanding our analysis to the macroevolutionary level, we use a database of over 1-million occurrence records and spatiotemporally-matched temperature data from 49 Swedish Odonate species to infer macroevolutionary dynamics of phenology. Phenological plasticity was more closely aligned with adaptation for species that have recently colonized northern latitudes, with more phenological mismatch at lower latitudes. Our results show that phenological plasticity plays a key role in microevolutionary dynamics within a single species, and such plasticity may have facilitated post-Pleistocene range expansion in this insect clade.

## Introduction

In many organism, life is characterized by dramatic life-history transitions between discrete stages that correspond to the unique demands of resource acquisition versus reproduction. For organisms with complex lifecycles, these life history transitions span disparate ecological niches, such as aquatic versus terrestrial environments, that demand irreversible metamorphosis to achieve such extreme ontogenetic niche shifts (Wilbur 1980; Werner & Gilliam 1984). The timing of such life history transitions, or phenology, is particularly crucial for organisms with complex life cycles that undergo their metamorphosis in seasonally variable environments. This is because success during a given life stage must depend not only on biotic factors such as predation, competition and seasonally-changing resources, but also on aspects of the abiotic environment, such as temperature, that vary across both time (generations) and space (between populations) (Hassal *et al.* 2007; Angilletta 2009; Plard *et al.* 2014; Visser *et al.* 2015; Bonamour *et al.* 2020). Thus, phenology is expected to be under net-stabilizing selection (Villemereuil *et al.* 2020) with an optimum that is shaped by various counteracting selection pressures that are in turn influenced by multiple biotic and abiotic environmental variables, including temperature, precipitation, competition and resource availability.

The location of phenological optima may fluctuate randomly in response to environmental stochasticity across years in seasonally variable environments (Hadfield 2016). Concomitantly, such fitness optima may show directional change and track advancing temperatures and resource abundances associated with anthropogenic climate change (Brashaw & Holzapfel 2001; Bradshaw & Holzapfel 2006; Parmesan 2006; Anderson *et al.* 2012; Thackeray & al. 2016). In both scenarios, phenological plasticity (phenotypic plasticity in the timing of life history transitions) is expected to be a target of natural selection (Chevin *et al.* 2010; Chevin & Lande 2015). Individuals that adaptively alter their developmental trajectory in response to available environmental cues regarding the conditions occurring during or after the life history transition (Whiteman 1994), will undergo metamorphosis to closely match the phenological optimum and will therefore have a selective advantage (Chevin *et al.* 2010). Phenotypic plasticity in phenology is therefore an expected key evolutionary adaptation to seasonally variable environments (Levins 1968). This classical hypothesis has obtained qualitative support in a number of studies across a range of taxa that all suggest a key role for plasticity in explaining between- and within population differences in phenological traits (Charmantier *et al.* 2008; Boutin & Lane 2014; Crozier & Hutchings 2014; Franks *et al.* 2014; Nashoba & Kono 2020; Stamp & Hadfield 2020). Only a few of these studies includeorganisms with complex life cycles (Phillimore *et al.* 2010; Stoks *et al.* 2014; Urban *et al.* 2014; Buckley & Kingsolver 2019).

Recent work has recast the concepts of plasticity, fitness, and environmental variation in terms of estimable quantitative genetic parameters that together describe plasticity’s potential contribution to adaptive evolution. In this framework (Figure 1) plasticity, termed *b*, and the environmental dependence of selection, termed *B*, jointly determine the extent that phenotypic plasticity contributes to local adaptation (Chevin *et al.* 2010). The strength of plasticity *b* describes the relationship between expressed population mean phenology and the average environment experienced by a population. This parameter (*b*) represents the population-mean reaction norm. The environmental dependence of selection, *B,* describes the relationship between the optimum phenology and the mean environment, and so represents the degree to which natural selection changes as a function of environmental variation. Importantly, estimation and comparison of these parameters can provide insight into plasticity’s role in adaptation (ability to track the optimum) to spatio-temporal environmental variation (Hadfield 2016). New statistical and analytical approaches have allowed researchers to leverage observational individual occurrence records to estimate these parameters indirectly, via space-for-time substitution (Phillimore *et al.* 2010). A growing body of studies have employed these or similar time-series based approaches (Phillimore *et al.* 2010; Crozier *et al.* 2011; Phillimore *et al.* 2012; Phillimore *et al.* 2016; del Mar Delgado *et al.* 2020; D. & Wilson 2021). This work (see also Villemereuil *et al.* 2020) indicates that plasticity can explain a substantial fraction of spatio-temporal variation in phenology across natural populations in the wild.

**Figure 1.**
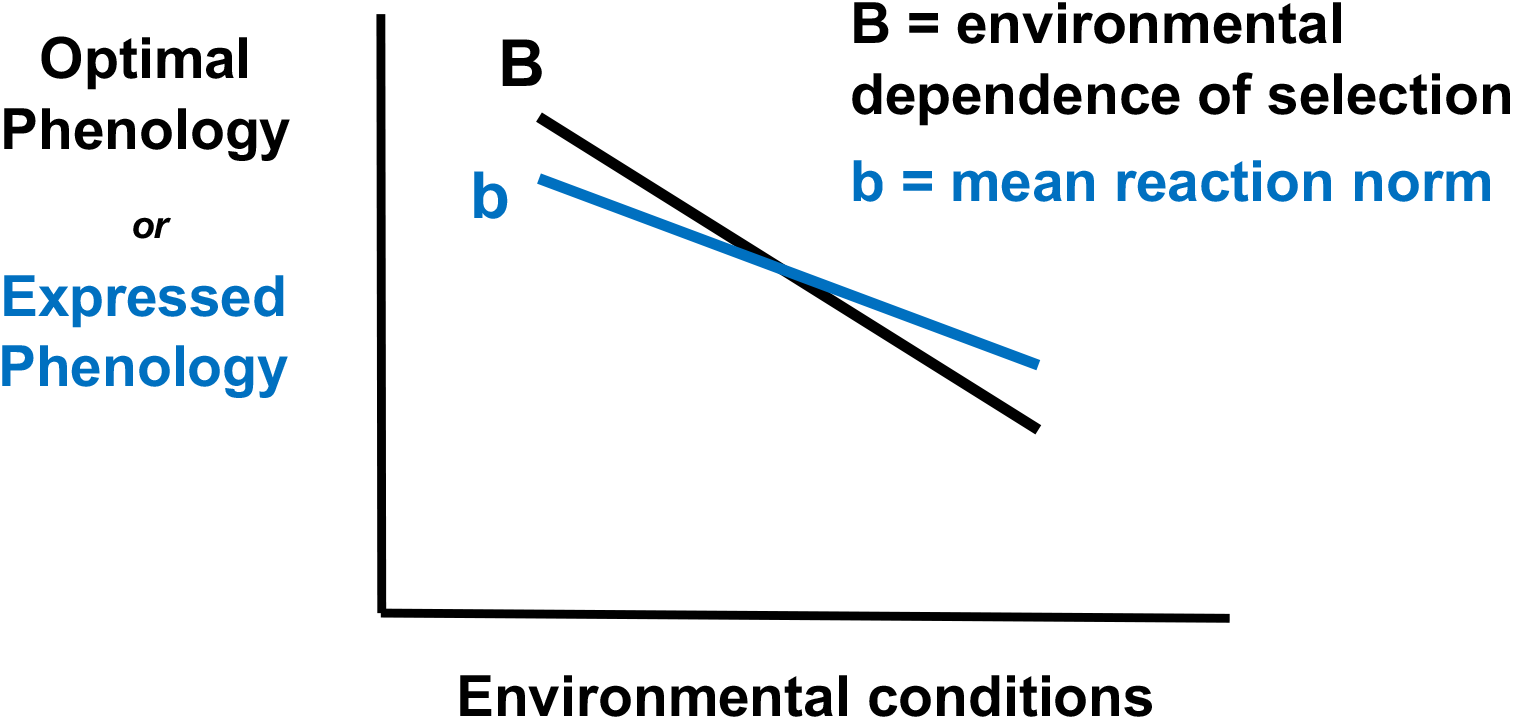
The environmental dependence of selection and the mean reaction norm jointly determine the role that phenotypic plasticity plays in adaptation. The x-axis represents an environmental variable important for individual fitness; the y-axis represents phenology, the timing of a key life history transition. The slope of the regression of the optimal phenology on environmental variation, B, represents the degree to which selection on phenology depends on the environment, shown in black. The degree to which expressed population mean phenology depends on environmental variation is represented by the population mean reaction norm, b, shown in blue. The degree to which B and b align determines the potential contribution of phenotypic plasticity to adaptation to environmental variation. Importantly, plasticity b is adaptive when it moves a population closer to the optimum than it would otherwise be following an environmental shift.

It is, however, still unclear whether and to what extent plastic variation in phenology matches our expectations for plasticity’s role in evolution. In particular, two critical open questions on phenological plasticity’s role in evolution remain unanswered: 1) does phenotypic plasticity correspond to environmental-dependence of phenological optima at the microevolutionary scale, thus amplifying or accelerating adaptation? and 2) is the role of plasticity in adaptation greater for lineages inhabiting variable and/or recently colonized environments? Answering the first question requires long-term data on individual fitness in the wild. Answering the second requires comparative analysis of multiple species or lineages that vary in ecology yet which share a similar and conserved complex life cycle.

Here we combine three independent, large-scale datasets from the insect order Odonata (damselflies and dragonflies). Odonates are semi-aquatic insects with a phylogenetically conserved complex life cycle characterized by metamorphosis from aquatic nymph (henceforth, ‘larvae’) to aerial reproductive adult (Stoks & Córdoba-Aguilar 2012). Our aim was to address the above two related questions on phenological plasticity’s role in adaptation at short and deep timescales.

### Methods – Data Collection

#### Long term study of Ischnura elegans

We analyzed data on individual phenology and fitness in a set of populations (approximately 16) of the damselfly *Ischnura elegans* in province of Skåne, southern Sweden, for 18 years (2000-2017). Each year, these populations were surveyed in a standardized approach during the reproductive season of *I. elegans*. All captured individuals were sexed and their copulation status (mated or not) was recorded. In this study, we used data on one major male fitness component: male mating success (a binomially distributed variable) to estimate the strength of selection on phenology through mating success.

#### Laboratory experiment on thermal plasticity in development time

We reared larvae of individual *I. elegans* from the egg stage, through hatching and metamorphosis to adults, under two different larval temperature environments: 20 °C and 24 °C. Maternal egg clutches (full sib clutches) were obtained from females caught *in copula* in the field in the general population survey described above. Individual eggs and larvae from a total of 101 full-sib clutches were reared in a split-clutch GxE design across both environments.

#### Occurrence records of Swedish Odonata

We obtained public, spatiotemporally referenced observational records of adult Swedish odonates identified to species level from the Swedish Species Observation System (Artportalen: https://www.artportalen.se/). We used data from a ten year period from 2006-2015, the range of years for which we had accurate temperature data and for which substantial observational records were available in this public database. We obtained spatiotemporally matched estimates of spring mean surface temperature from the publicly accessible NOAA Global Forecast System Analysis (GFS ANL) database (https://www.ncdc.noaa.gov/data-access/model-data/model-datasets/global-forcast-system-gfs). We obtained the mean surface temperature from corresponding lat-long grid cell in the GFS ANL database for two time windows: the first 120 days of the year and the month of April. The first time window corresponds to a substantial period of pre-metamorphosis larval development and covers the last instar for all the species in our database, and so represents a long-term average temperature environment experienced during a key stage of development. The second time window represents a shorter time slice from late spring, just prior to the flight period of many early season species. Finally, we note that the scale and purpose of our analysis precludes use of a “sliding window” approach; Our interest is to compare developmental-temperature reaction norm variation across species (and so any such comparison requires use of similar or the same environmental cues), rather than to identify a specific time slice of spring temperature variation that is most important for a single species.

### Methods – Statistical Analysis

#### Long term field study of Ischnura elegans

To infer selection on phenology, we modify the Lande-Arnold (Lande & Arnold 1983) equation to include temperature dependence in the form of directional selection on phenology:

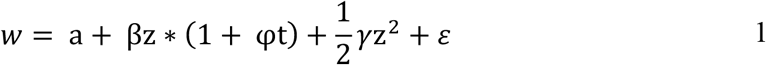

Where 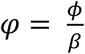, *ϕ* is an interaction term as estimable in a regression model. Setting the first derivative of equation 1 with respect to *z* to zero and solving yields the stationary point:

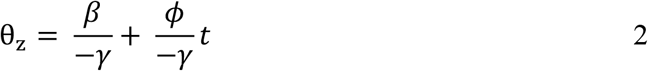

where θ_7_ is the optimum phenology (if selection is stabilizing, otherwise it is a fitness minimum). The slope of the relationship between temperature and θ_z_ is 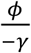, and represents a direct estimate of the temperature dependence of selection (*B*).

#### Laboratory temperature experiment: raising damselfly families

We used a mixed effects model to estimate the relationship between thermal developmental environment of larvae (two temperature treatments) and emergence time from our laboratory experiment. Our model featured emergence day as the response variable, temperature treatment as a fixed effect, and random intercepts for tank and maternal full-sib family

#### Occurrence records of Swedish Odonata and phylogenetic comparative analyses

We used a bivariate mixed modelling approach (Phillimore *et al.* 2010; Phillimore *et al.* 2012) to infer interspecific variation in thermal reaction norms in phenology from our observational records of Swedish Odonata combined with paired estimates of temperature in the first 120 days of the year. We fit the bivariate model

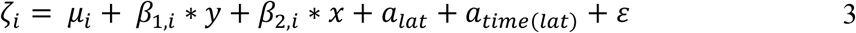

where *ζ*_i_ is the response vector of individual values of two variables: *i* = observed phenology or spring temperature. *β* is a fixed effect regression coefficient between continuous latitude (*y*) or year (*x*) and variable *i*. Two nested random effects are included in this model: *a_lat_* is a random effect describing (co)variation across binned latitude, and *a_time(lat)_* is a random effect describing co(variation) across binned annual seasons within latitudinal grids. For each *a* and the residual *ε* we estimated the full 2×2 unstructured covariance matrix 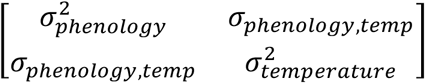, for a total of three 2×2 random effect covariance matrices. We focus on analysis of latitudinal spatial variation because Sweden spans a far greater latitudinal than longitudinal range, and corresponding species distributions within Sweden reflect substantial latitudinal span but little longitudinal span.

From this model we can interpret the ratio of the fixed effect parameter estimates, particularly 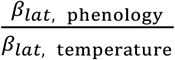 as approximating the environmental dependence of the optimum, *B* (Hadfield 2016). From the random effects, we can interpret *b* ∼ *σ*_phenology, temperature_/ 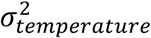 where *σ* and *σ*^2^ are estimated across time points within binned latitude (*a_time(lat)_*). This bivariate model was fit separately for each species in our dataset. We then used the corresponding estimates of *B* and *b* in phylogenetic mixed models (Hadfield & Nakagawa 2010) to explore the relationship between species mean latitude, species mean phenology, and the strength and form of inferred phenological plasticity. Our models accommodated uncertainty in our estimates of *B* and *b*.

## Results

We found evidence of significant phenological advance during the 18 year study period of *I. elegans* in the field (F_1, 47526_ = 1231.38, P <0.0001, Figure 2B). Although there was significant among-year heterogeneity, on average the emergence date has advanced by .58 days (SE = 0.017) per year, corresponding to a predicted total advance of 10.44 days across the entire 18-year period.

**Figure 2.**
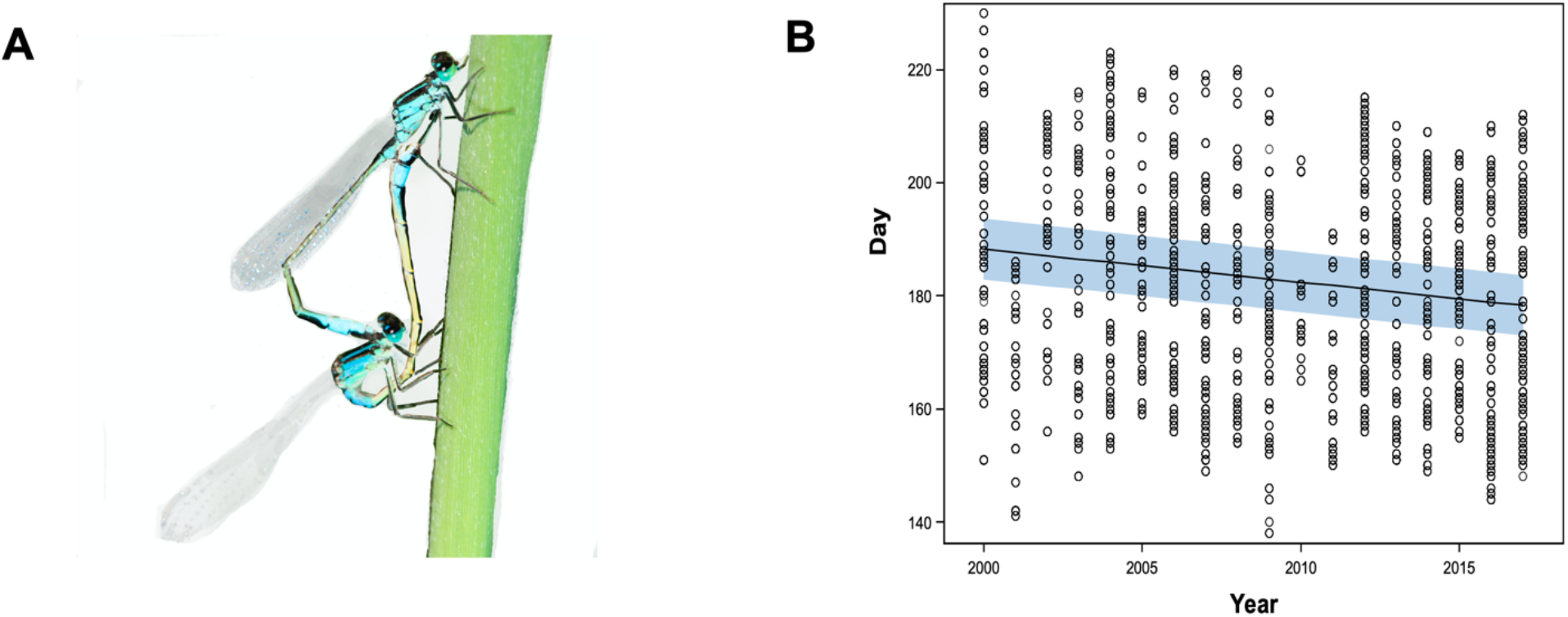
Reproductive advance in *I. elegans* in Southern Sweden. *Ischnura elegans*, a common European damselfly, exhibits an aerial reproductive adult stage (**A**) that follows a prolonged aquatic larval growth stage. Long term study of a set of populations in southern Sweden over 18 years indicates a significant advance in the timing of the aerial adult phase (**B**). Conclusions were unaffected by exclusion of data from the year 2000. Blue shading in B shows 95% confidence limit of the prediction.

Using data on male mating success (N = 21,384 individuals for which we also had annual temperature data, consisting of 5,487 mated males and 15,897 non-mated males), we find evidence of statistically significant but annually variable (Figure 3B) sexual selection on phenology. Overall, we found evidence of significant stabilizing selection on phenology (estimates of parameters in equation 1, in units of raw days: *γ* = −0.00016, SE = 0.000072, 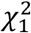 = 4.51, P = 0.0336; β = 0.001308, SE = 0.000972, 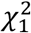 = 1.18, P = 0.2; binomial glmm: *γ* 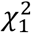 = 6.99, P = 0.0177; β 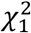 = 2.16, P = 0.16), along with a significant temperature-dependence of selection on phenology (*ϕ* = −0.00156, SE = 0.000593, 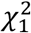 = 6.90, P = 0.0086; binomial glmm: 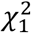 = 5.02, P = 0.0251). Fitting a simplified model with directional selection only and random variation among years to variance-standardized data indicates standardized directional selection gradients varied substantially among seasons (ΔAIC = 265, overall estimate β = 0.027, SE = 0.021; see figure S1). The resulting estimate of the relationship between θ_z_ and spring mean temperature indicates a temporal advance in the optimum phenology with increasing temperature (Figure 3C). Our point estimate of *B*, the slope of the relationship between temperature and θ_z_, given by the ratio of the parameter estimate 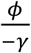, indicates an advance of 10.24 days (Bootstrapped 95% CI: −66.7995 – −0.59705) in the phenological optimum for each degree Celsius of spring warming (Figure 3C). We also found evidence of negative *B* using an independent dataset of 1695 observations from the Artportalen database when we inferred B indirectly from interspecific data on phenology, although this indirect estimate was significantly shallower than direct estimate we obtained from this long-term population study (Supplemental material, Figure S2).

**Figure 3.**
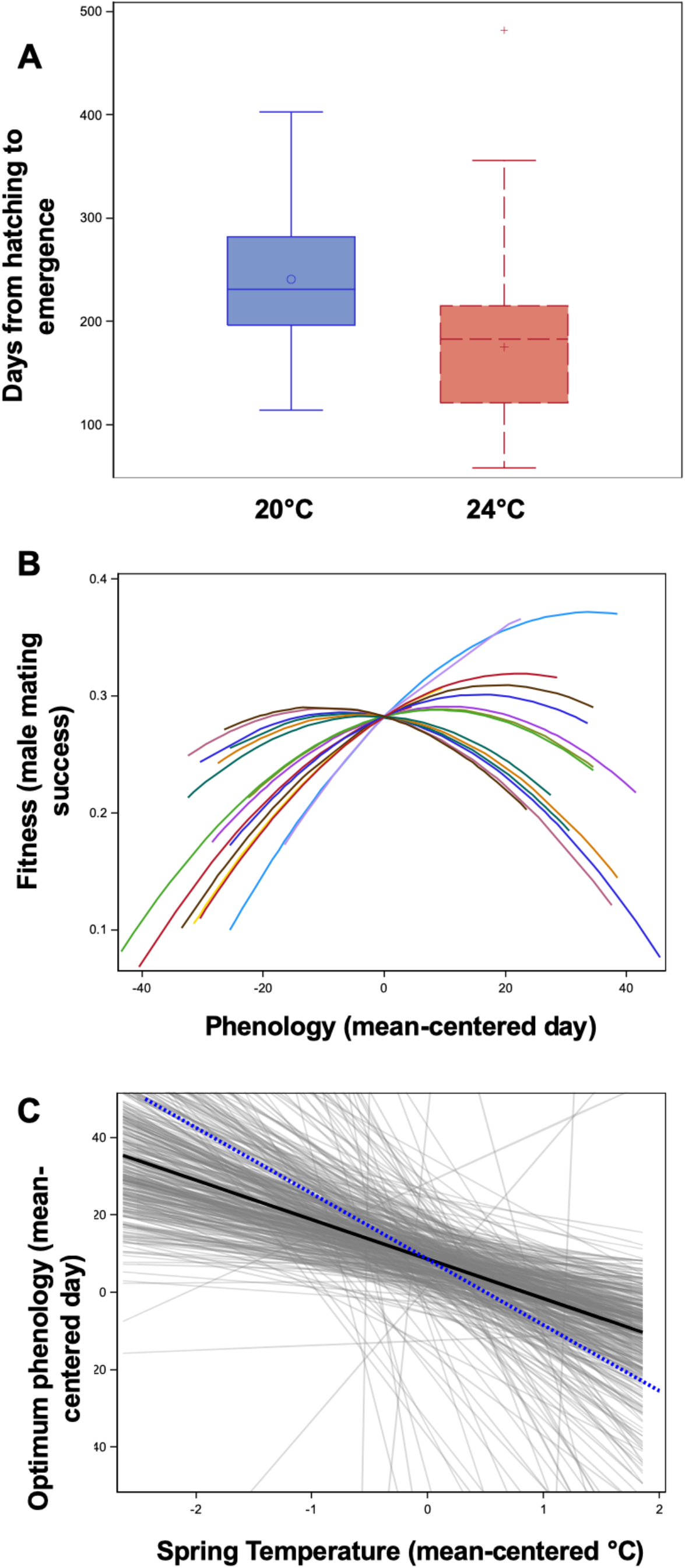
Temperature dependent selection and plasticity are aligned in *I. elegans*. In a laboratory experiment, increased larval developmental temperature was associated with accelerated time to metamorphosis as adults. **A**. Yaxis shows days from hatching to emergence. In a long term study of *I. elegans* in the wild, selection on phenology (measured as male mating success) varied across years but overall, selection was net-stabilizing, **B**, with optima that vary slightly across years (lines show model fits from the fixed effects across years; each color represents a unique year). The estimated optimum was negative temperature dependent, **C**.Panel C shows the linear relationship between θ_z_ and spring mean temperature (equation 2 in text); black line shows original ML estimates, grey lines show 400 samples from a multivariate normal distribution centered at the ML estimates with covariance equal to the covariance matrix of fixed effected from the fitted model (i.e. a parametric bootstrap). Dashed blue line shows the estimated reaction norm from the plasticity experiment (see panel A).

We obtained individual and family emergence time records from 521 individuals from 101 full-sib genetic families across two temperature treatments in our laboratory experiment. We found significant acceleration of development in response to our experimental manipulation of temperature (F_1, 367_ = 65.61, P <0.0001, Figure 3A), with an estimated reaction norm of −17.38 (SE = 2.15) days per 1 °C warming of the developmental environment. This laboratory estimate therefore corresponds in sign, and to some extent in magnitude, with our estimate of the environmental dependence of θ_z_ estimated from the field data in the natural populations of *I. elegans* (Figure 3C). We found statistically significant broad-sense heritability in the timing emergence (*H*^2^ = 0.24, 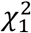= 17.54, P < 0.0001; SE: 0.078, 95% CI: 0.079-0.39; V_a_ = 993 SE: 245, V_p_ = 4156 SE = 326; CV_A_ = 5.3%). Our finding of a negative thermal reaction norm in larval development time that aligns with the environmental dependence of selection on phenology was supported by an independent, indirect inference of *b* using observational data (although the magnitude of these estimates differs substantially; Figure S2).

We analyzed paired temperature and phenology data from 1,027,952 records from 49 species of Swedish Odonates (see Table S1). We explored two temperature windows as predictors of phenological variation: the mean temperature of the first 120 days of the year, and the mean April temperature. We found substantial support for the 120 day window, as these models consistently fit better than the April mean models for nearly all species (Figure S3). We then kept the 120 day average models for downstream analysis.

Reaction norm estimates varied across species, although most estimates where within the range of ± five days per °C of spring temperature variation (Figure 4A, Table S1). Notably, there were a mix of both positive and negative reaction norm estimates and many species that showed positive reaction norm slopes (Fig. 4). These analyses show that temperature during the last instar larval stage can either advance or delay emergence, depending on species, and some species were relatively insensitive to spring temperature (Fig. 4A). On average, the estimated thermal reaction norms explained 1.4% of the all variation in phenology (Table S1). In comparison to our estimates of b, estimates of B exhibited substantial sampling variance (Table S1).

**Figure 4.**
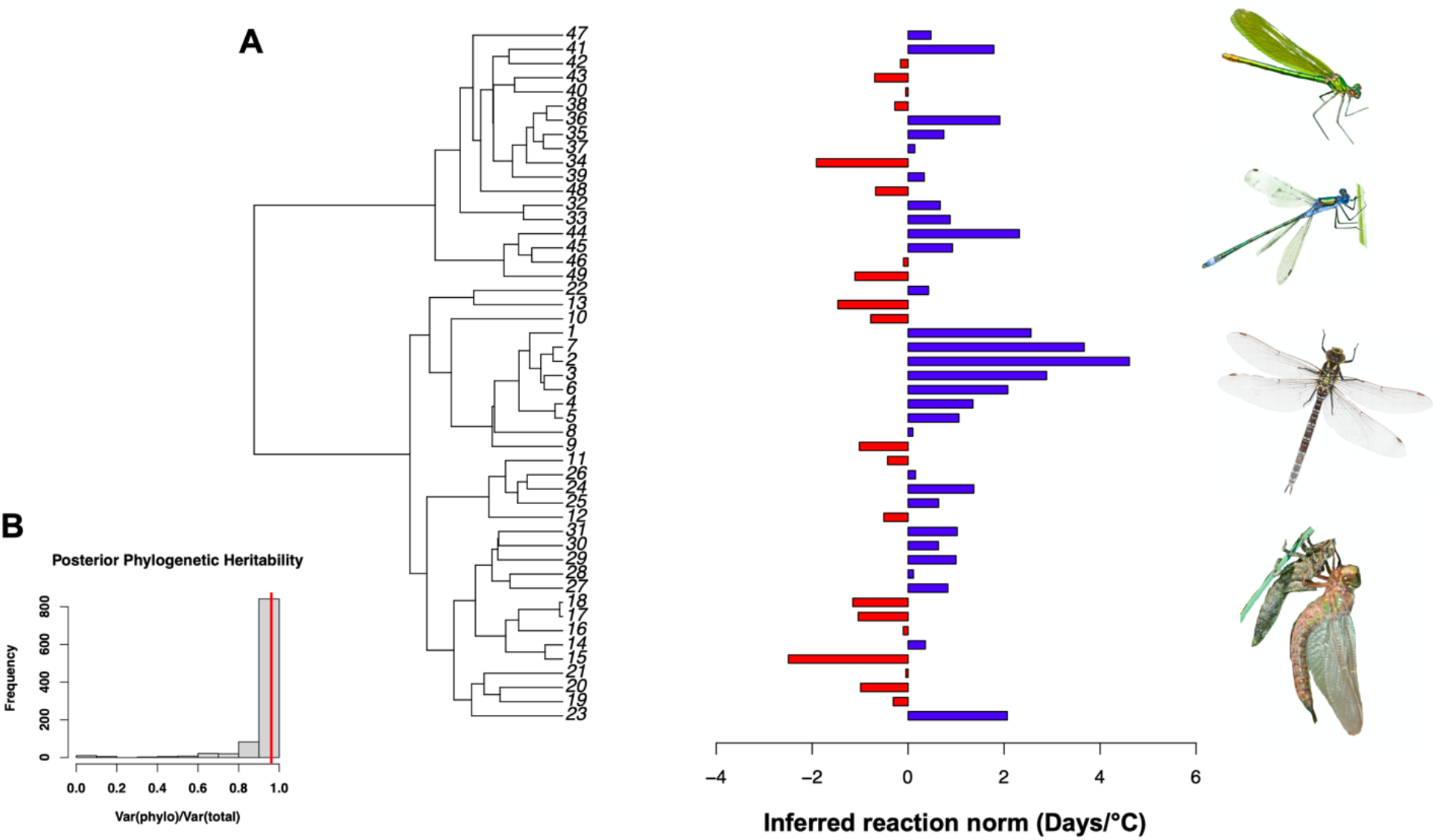
Inferred temperature reaction norms for phenology across Swedish Odonates. **A,** Estimated reaction norms for temperature dependent phenology show variation among 49 damselfly and dragonflyspecies, although most are relatively shallow and the phylogenetic signal is high and statistically significant. Species images (photos by S. De Lisle, top down): *Calopteryx splendens* (ID 32), *Lestes dryas* (ID 44), *Aeshna cyanea* (ID 1), *Brachytron praetense* (ID 9). Full species names for each ID number are provided in Table S1. Panel **B** shows the posterior distribution and point estimate (red line) for the phylogenetic heritability of the phenological reaction norm.

We found evidence of a significant phylogenetic signal in the phenological reaction norm (Phylogenetic *h*^2^ = 0.96 ΔDIC = 239, posterior distribution in Figure 4B), revealing that closely related species have similar magnitude of thermal plasticity in phenology (Figure 4). Estimated reaction norms were correlated with mean phenology across species, with early season species exhibiting more negative reaction norms and late season species exhibiting positive reaction norms (posterior mean coefficient: 0.03192, 95% CI: 0.01915 – 0.04435, pMCMC < 0.001, Figure 5A). We also found a significant negative relationship between reaction norm slope and latitude in a phylogenetic mixed model with latitude as a predictor of estimated reaction norm (posterior mean coefficient: 0.40298, 95% CI: 0.03198 – 0.78868, pMCMC = 0.036, Figure 5B).

**Figure 5.**
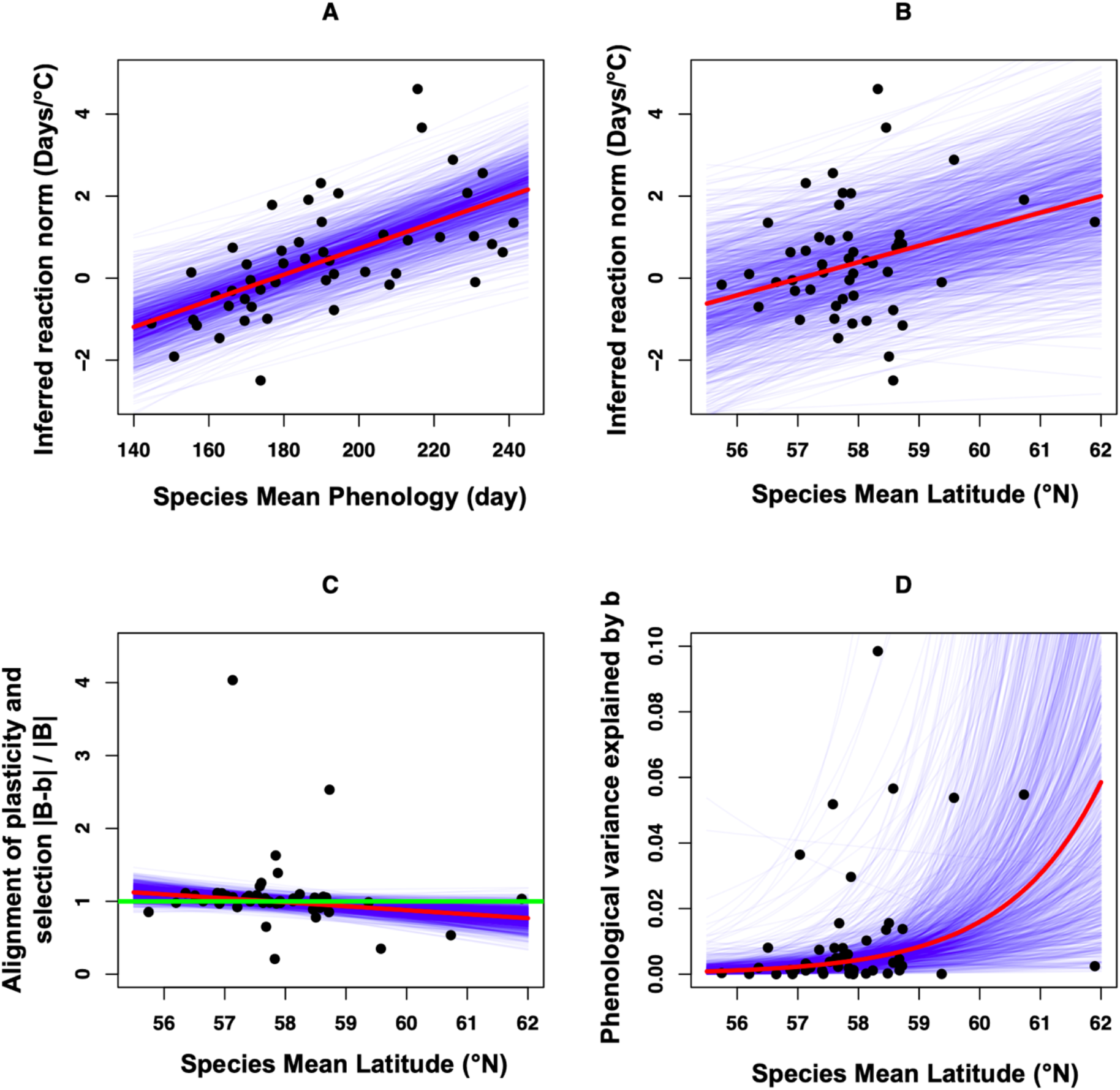
Macroevolutionary dynamics of phenological plasticity. Early-emerging odonate species have more negative thermal reaction norms than late-emerging species. Species that emerge late in the season instead tend to exhibit positive reaction norms to winter/spring temperature, **A**. There is evidence that negative reaction norms, reflecting accelerated development in response to warm temperatures, evolve in southern latitudes **B**. Across species, the alignment between inferred reaction norm slope, b, and the inferred environmental dependence of the optimum phenology B, becomes greater (smaller absolute difference scaled by the magnitude of B) for species with Northern geographic ranges **C**. Green line in C equals unity; values of |B-b|/|B| < 1 indicate adaptive plasticity and values > 1 maladaptive plasticity. Inferred plasticity explains a greater proportion of the detrended variation in phenology in Northern species, Panel **D**. Together these data suggest that reaction norms have diversified at the macroevolutionary scale and that this diversity may have played a an important role during Odonate range expansion into northern latitudes. In all panels, red lines show posterior mean model fits, blue lines show the posterior distribution. All models accounted for phylogenetic relationships and uncertainty in the species-specific estimates of the *B* and *b*. In panel **C** an extreme single outlier (*Aeshna grandis*) is not shown in the figure, but inclusion/exclusion of this species had no effect on general conclusions (see text).

The alignment between the estimated phenological reaction norms, b, and the estimated environmental dependence of selection, B, scaled by the magnitude of B (i.e., |B-b|/|B|), varied significantly with species mean latitude (posterior mean coefficient: −0.054454, 95% CI: - 0.115952 – −0.004226, pMCMC = 0.048; with outlier dropped, posterior mean coefficient: - 0.055290, 95% CI: −0.105413 – 0.004944, pMCMC = 0.046; Figure 5C). When |B-b|/|B| is less than unity plasticity can be considered adaptive; when above unity plasticity can be described as maladaptive (Phillimore *et al.* 2016). Species at northern latitudes had thermal reaction norms that were more closely aligned with their estimated environmental dependence of phenological optima, resulting in adaptive plasticity at northern latitudes (Figure 5C). Finally, plasticity *b* explained a greater amount of detrended variation in phenology (calculated as 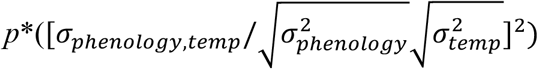 were the term in parentheses is at the level of time(lat) and *p* is the proportion of the total variance in phenology that is explained by the time(lat) level) in Northern than in Southern species (posterior mean coefficient: −0.6489, 95% CI: - 1.1821 – - 0.1102, pMCMC = 0.018, Figure 5D).

## Discussion

Here we combine three independent datasets, together representing well over 1 million individual insect records, to test two related questions on the role of phenotypic plasticity in the evolution of phenology. First, we demonstrate pronounced recent phenological advance in a single damselfly species, *Ischnura elegans*, using a long term study of individuals in the wild. We find that thermal plasticity in developmental time in the laboratory is substantial, corresponding in sign and to some degree in magnitude with the relationship between temperature and the optimum phenology of the same species estimated in the field. This suggests an important role for thermal plasticity in adaptation to seasonal environmental variability, and also suggests plasticity could contribute to advancing phenology that corresponds to recent anthropogenically-caused climate change in Sweden (SMHI 2015). At the macroevolutionary scale, we used a large interspecific database of individual emergence records of 49 species of odonates spanning the extensive latitudinal gradient of Sweden. Combining these occurrence records with spatiotemporally-matched estimates of spring mean temperature, we found that species-specific mean phenological reaction norms are more positive and more closely aligned with indirect inferences of the environmental dependence of the optimum for northern than for southern species. We also find that plasticity contributes more to variation in phenology at Northern latitudes. Our combined intra- and interspecific results indicate that temperature-dependent plasticity in phenology underlies both microevolutionary adaptation to contemporary climate change as well as the macroevolutionary expansion into harsh high-latitude climatic conditions that took place when species colonized these areas after the last Ice Age in Fennoscandia, about 10,000 years before present.

We found a phenological advance in a single species (*Ischnura elegans*) of 0.58 days/year, almost six days per decade or 10.44 days over the entire 18-year study period (Fig. 1). This is a dramatic phenological advance during the first two decades of the century in Sweden. In fact, this is almost twice as large advancement of phenology as was documented previously for *I. elegans* in a study of British Odonata for the period 1960-2004 (0.58 vs. 0.33; Hassall *et al.* 2007). Thus, although our findings of phenological advance are consistent with previous study in Britain, the new results presented here suggest temperature-dependent phenological advance in odonates may be more extreme than previously appreciated.

We find some correspondence between point estimates of the environmental dependence of selection in *Ischnura elegans* and the population mean reaction norm, using three independent datasets and approaches. Our direct estimate of *B* using data on individual fitness from the long-term study on *I. elegans* and our indirect estimates using a public database of individual occurrence records both suggest that the optimum emergence time advances substantially for every centigrade degree of spring warming in *I. elegans*. Our direct estimate of the phenological reaction norm matches with our direct estimate of the environmental dependence of selection, and the correspondence is also found using a bivariate mixed model approach to indirectly estimate these parameters (see Supplemental Material). Although absolute values of *B* and *b* differ substantially across these two approaches (direct vs indirect), the qualitative conclusion – that phenotypic plasticity can explain the close match between local optima set by environmental variation – is similar across these different approaches and datasets. One the one hand, this correspondence in qualitative interpretation (alignment of *B* and *b*) is encouraging and suggests that our space-for-time substitution approach to estimating environmental dependence of optimal phenologies may be robust to inevitable violation of the assumptions (e.g., local adaptation and limited dispersal) that underlie the theory behind the method (Hadfield 2016). On the other hand, we found a significant disparity in the magnitude of both *B* and *b* using these different approaches; our estimates of *B* and *b* were significantly steeper in our direct assessment (Figure S2). A likely explanation for this is that our direct estimate of these parameters was focused on a more limited section of the geographic range, compared to our indirect estimate using data from *I. elegans* across the latitudinal breadth of Sweden. In another comparison between these types of approaches, Phillimore et al. (Phillimore *et al.* 2016) found relatively close correspondence between different estimates of *B* in great tits (see also van Benthem *et al.* 2017 for a comparison of other related approaches). The results presented here echo past suggestions (Phillimore *et al.* 2016) that further validation of indirect estimates of *B* is warranted.

Our laboratory breeding experiment revealed a strong effect of temperature at larval stage on development and emergence time, with an advance of 17 days per degree Celsius. The fact that this estimate of the mean reaction norm was steeper than our estimate of the environmental-dependence of selection might suggest ‘hyper plasticity’ (Stamp & Hadfield 2020) wherein plastic responses have the potential to exceed fluctuations in the optimum (see Radersma et al. 2020). Nevertheless, our laboratory experiment indicates that the strength of temperature-dependent plasticity is qualitatively similar to our estimate of the environmental dependence of selection and is broadly aligned with the direction of temporal phenological advance in response to rising temperatures in natural populations (Fig. 3C). Indeed, the estimated population reaction norm in *I. elegans* suggests that an adaptive fit between the thermal optimum and mean phenology could be explained entirely by phenotypic plasticity. This indicates that *I. elegans* may already be well-adapted to cope with changing climates and increasing temperature compared to many other animal taxa, such as birds (Radchuk *et al.* 2019). Previous studies on gene expression of heat- and cold shock proteins in *I. elegans* adults originating from different parts of the geographic range indicate that adaptive thermal plasticity plays an important role in the northward range expansion of this species (Lancaster *et al.* 2015; Lancaster *et al.* 2016), in addition to genetic adaptation (Dudaniec *et al.* 2018).

Our laboratory experiment also revealed that development time was modestly heritable, with a point estimate of *H*^2^ = 0.24. This is considerably lower than a previous estimate of the heritability of development time in this species (0.52), estimated in a single thermal environment (Abbott & Svensson 2010). The modest level of heritability of development that we document here would require a persistent variance-standardized directional selection gradient of approximately *β* = -.13 to result in the observed annual phenological advance of 0.58 days/year (approximately .03 within-year standard deviations). This would be relatively strong selection compared to available evidence from field studies on phenotypic selection (Kingsolver *et al.* 2001). Although our results and analyses do indeed suggest that selection can be strong, we also find little evidence of persistent directional selection of the magnitude required to produce such a dramatic response. In fact, across all years, net selection on phenology is actually slightly positive rather than negative as would be expected if earlier phenology was consistently favored (Fig. S1). Therefore, it is unlikely that genetic evolution alone can entirely explain the strong phenological advance of more than 10 days across the entire 18-year period, given the substantial among-year variation (Figure 2) of *I. elegans* in southern Sweden. Instead, our analyses suggest that phenotypic plasticity may have an important role in driving both inter-annual variation in phenology and the progressively earlier flight seasons that we have documented here. These data suggest that *I. elegans* can track changing phenological optima through plasticity, which will in effect reduce or counteract directional selection for earlier phenology, a conclusion that was also reached in a recent study of 21 bird and mammal species with long-term data (Villemereuil *et al.* 2020). More generally, various forms of phenotypic plasticity have been proposed to “buffer” organisms against external stress from temperature extremes (Huey *et al.* 2003).

We found significant, phylogenetically-structured among-species variation in inferred reaction norms in an analysis that explicitly accounted for uncertainty in these species-specific estimates. Our point estimate indicates a phylogenetic heritability of near unity, indicating that closely related species share similar phenological responses to temperature variation. The significant phylogenetic signal in the reaction norm estimates is consistent with the idea that species differences in ecology and physiology play important roles in generating reaction norm variation across this clade, most likely because closely related species show a high degree of niche conservatism (Wiens & Graham 2005; Svensson 2012; Wellenreuther *et al.* 2012). Our findings add to a growing body of evidence from other taxa (Davis *et al.* 2010; Davies *et al.* 2013; Rafferty & Nabity 2017), indicating strong phylogenetic signal in phenological response to climatic change.

Theory suggests that the fitness benefit of phenological plasticity depends upon the existence of environmental cues that accurately predict the conditions that determine fitness in the next life stage (Chevin & Lande 2015). Here we have focused on spring mean temperature as a potential environmental cue (or correlate thereof). This is justified by substantial empirical evidence linking temperature and corresponding thermal tolerance to various aspects of adult fitness in insects in general (Punzalan *et al.* 2008; Garcia-Roa *et al.* 2020), and our study species *I. elegans* in particular (Lancaster *et al.* 2015; Lancaster *et al.* 2016; Dudaniec *et al.* 2018; Svensson *et al.* 2020b). Moreover, our laboratory experiment confirms that larval developmental temperature is a predictor of emergence time into the transition to the adult aerial life-stage. Across all our interspecific dataset, all but one species in our comparative analysis over-winter as either aquatic larvae or eggs, and so are exposed to spring temperatures at a similar developmental stage. The only exception to this rule is the winter damselfly *Sympecma fusca*, which overwinters as an adult in vegetation refugia. Consistent with this, *S. fusca* exhibited one of the most extreme negative estimate of *B* (Table S1), as well as a negative estimate of *b* (Fig. 4A). This fits with previous research showing that temperature-phenology reaction norms tend to be strongest in insects that over winter as adults (Forrest 2016). Across all the species included in this study, other environmental cues are also likely to be important for determining developmental rate; including aspects of the biotic environment, such as density of con- or heterospecific competitors, resource availability, and photoperiod. These unknown additional cues could explain much of the residual variance in phenology. Nevertheless, our focus on temperature is relevant in light of anthropogenic climate change (Parmesan 2006) and the general consensus that temperature is a strong determinant of fitness in ectotherms (Angiletta 2009).

The degree of alignment between the phenological reaction norms and environmental dependence of selection, and the contribution of plasticity to variation in phenology, changed across the latitudinal range of these 49 species in Sweden (Fig. 5). This indicates that adaptive phenological plasticity has played an important role in the northern range expansion of this clade. Insect colonization of the Scandinavian peninsula occurred relatively recently following retreat of the Fenno-Scandian ice sheet, which covered the region from the last glacial maximum until its recession at the end of the Pleistocene approximately 10,000 years ago (Hewitt 1999, 2000). Thus, odonate species that currently occur in northern Sweden have established themselves in this region more recently than the ones in southern Sweden. Our findings of stronger evidence of adaptive plasticity in northern species in our phylogenetic comparative analysis (Fig. 5) is consistent with theoretical predictions (Lande 2009; Chevin & Lande 2011) that plasticity might be particularly important in adaption to novel and variable environments, such as near range limits (Lancaster *et al.* 2015; Lancaster *et al.* 2016). Conversely, we found that southern species currently show some degree of non-adaptive or even maladaptive plasticity in relation to temperature as they show values of scaled |B-b| greater than unity (Fig. 5C). A caveat here is that our estimates of *B* (both direct and indirect) exhibit substantial sampling variance (Table S1; Results, Figure 3C), due in large part to ratio effects that amplify effects of latitudinal regression coefficients / selection coefficients that compose the indirect/direct estimate of *B*, respectively. Even our direct estimate of *B* from *I. elegans*, inferred using a large dataset of individual component fitness measures, exhibited high sampling variance, which highlights the challenges of estimating and interpreting these parameters even in the presence of a large dataset.

Despite these challenges, our analyses are broadly consistent with recent research and indicate that phenotypic plasticity can be adaptive (Noble *et al.* 2019; Stamp & Hadfield 2020; Svensson *et al.* 2020a; Johansson *et al.* 2021), although we note that the much observed plasticity in our study is not necessarily adaptive. Interestingly, there was some degree of mis-alignment between *b* and *B* in the southern species *I. elegans* (Fig. 3C), which is broadly consistent with the greater mis-alignment in southern, compared to northern species at the macroevolutionary level (Fig. 5C). More generally, plasticity can be adaptive and selected against in unpredictable environments (Leung *et al.* 2020) and recent meta-analyses have documented that the degree of plasticity sometimes exceeds genetic differentiation, revealing non-adaptive or maladaptive plasticity (Radersma *et al.* 2020; Stamp & Hadfield 2020).

By combining multiple and independent datasets, we have revealed a potential role for phenotypic plasticity in explaining intra- and interspecific variation in phenology. Our combined intraspecific and phylogenetic-comparative analyses enabled us to evaluate the role of phenological plasticity in post-Pleistocene micro- and macroevolution of the temperature-dependence of insect phenology. Our work indicates that temperature-dependent phenological plasticity can at least partly track changing optima due to recent climate change, and that such adaptive phenological plasticity may also have facilitated historical range expansion into variable and extreme northern environments.

## Acknowledgements

We thank the many student interns that assisted with collection of the long-term *I. elegans* data, and especially Rachel Thomson and Tilly Pembury Smith who helped carry out the laboratory breeding experiment during 2017-2018. Deborah Goedert and Ally Phillimore provided critical feedback on the manuscript. Funding was provided by a postdoctoral scholarship grant to M.M. from The Emil Aaltonen Foundation, grants from the Royal Swedish Academy of Sciences and the Royal Physiographical Society of Lund to S.P.D., and research grants from The Swedish Research Council (VR; grant no. 2016-03356), Gyllenstiernska Krapperupstiftelsen (grant no. KR2018—0038), Lunds Djurskyddsfond, “Olle Engqvist Byggmästare Foundation”, John Templeton Foundation (grant no. 60501) and Stina Werners Foundation to E.I.S.

## Supplemental Material

### Detailed Methods-Data Collection

#### Long term study of Ischnura elegans

We analyzed data on individual phenology and fitness in a set of populations (approximately 16) of the damselfly *Ischnura elegans* in province of Skåne, southern Sweden, for 18 years (2000-2017). This set of populations spans an area of approximately 40×40 km. Each year, these populations were surveyed in a standardized approach during the reproductive season of *I. elegans*, beginning in late May when they start to emerge from the aquatic larval stage and continuing throughout the major part of the flying season until the end of July. Field work took place daily, except during days of low temperature (< 14 °C), heavy wind and rain, when adult damselflies are not active. Each population was visited at intervals between one and two weeks, within the lifespan limits for adult *I. elegans* and corresponding to approximately 5 sampling visits per site per year (Willink & Svensson 2017). Sampling was conducted for approximately 60 minutes at each site during a sampling session. See Willink and Svensson (2017) for more details about general field and laboratory work procedures.

*I. elegans* adults were captured in the field. Copulating pairs were kept separate from single-caught individuals in individual plastic containers. In the laboratory, all individuals were sexed and their copulation status (mated or not) was recorded. In this study, we used data on one major male fitness component: male mating success (a binomially distributed variable) to estimate the strength of selection on phenology through mating success. Our focus on male component fitness was due to two reasons: first, statistical power, in that we obtained far more estimates of male mating success across seasons, and the nonlinear, random-regression approaches (see below) to estimating variation in selection are data hungry. Second, our previous work has shown that this fitness measure is correlated with female fitness (fecundity) in these populations, corresponding to male and female phenotypic selection estimates that are significantly and positively correlated with each other across these populations (Gosden & Svensson 2008). This finding indicates that a broad patterns of selection acting through male component fitness are indeed reflective of patterns of selection acting through female fecundity in this system. We also note that female multiple mating is rare due to mate guarding in copula. Hence, male mating success is a reasonable proxy of mean population fitness as a whole. Our surveys yielded data on phenology (date of capture) from 47,615 individuals (28,978 males, 18,637 females) during this 18 year period (2000-2017). As the majority of individuals of *I. elegans* in these populations in southern Sweden are univoltine and emerge as adults after one year in the larval stage, these 18 years correspond to approximately the same number of generations.

#### Laboratory experiment on thermal plasticity in development time

We reared larvae of individual *I. elegans* from the egg stage, through hatching and metamorphosis to adults, under two different larval temperature environments: 20 °C and 24 °C. These temperatures were chosen for two reasons. First, they mimic typical and elevated summer conditions in Sweden, respectively, under a projected climatic shift of 4°C (De Block *et al.* 2013; Dinh *et al.* 2013). Second, past work in odonates has indicated that temperature differences on this scale can manifest significant affects on larval development (Dinh *et al.* 2013). Our choice of only two temperature treatments also precludes estimation of more complex, nonlinear reaction norms. Although this is a limitation of our design, we note that because statistical theory (Phillimore *et al.* 2010; Chevin *et al.* 2015) behind reaction norm inference at other levels of analysis in our study is limited to the case of linearity, we are most interested in the linear approximation of the reaction norm. Maternal egg clutches (full sib clutches) were obtained in 2017 from females caught *in copula* in the field in the general population survey described above. Females caught *in copula* were set up for oviposition in individual plastic cups at room temperature, and provided with a wet filter paper for egg laying surface. Two days later, the female was removed, the eggs were scanned and counted from the digital images. See Svensson and Abbott (2005) and Svensson et al. (2005) for further methodological details about laboratory procedures. Individual eggs and larvae from a total of 101 full-sib clutches were reared in a split-clutch GxE design across both environments. We started with 20 full siblings per environmental treatment for a total of approximately 4000 larvae across this rearing experiment. Upon hatching, individual larvae were transferred to plastic cups where they were housed individually in temperature-controlled water baths. Larvae were fed *Artemia* daily and kept at a constant midsummer light cycle (16:8). Towards the end of larval development and when the larvae reached the last instar, each cup received a wooden stick as an emergence aid and cups were covered with mesh to prevent escape of emerging adults. This experiment continued until all individuals had either died as larvae or emerged as adults. Of the 4000 larvae, 523 (13%) successfully emerged as adults, with recorded emergence time for 521 individuals. Although mortality in this experiment may appear nominally high, it is consistent with the high larval mortality observed in other odonates (Anholt 1994; Stoks & Córdoba-Aguilar 2012), and we note that over 500 adults recruited from 101 females corresponds to an absolute fitness (discrete time per capita growth rate) ≍ 5, an exceptionally high number. Larval development time was calculated as date of emergence as an adult minus the date of hatching, consistent with our previous procedures (Abbott & Svensson 2005).

#### Occurrence records of Swedish Odonata

We obtained public, spatiotemporally referenced observational records of adult Swedish odonates identified to species level from the Swedish Species Observation System (Artportalen: https://www.artportalen.se/). These data represent a combination of validated citizen science records, as well as records reported by professional scientists and naturalists throughout Sweden, and these data are also subsequently exported to the Global Biodiversity Information Facility (GBIF: https://www.gbif.org/). We used data from a ten year period from 2006-2015, the range of years for which we had accurate temperature data (see below) and for which substantial observational records were available in this public database. We excluded observations of larvae, as well as for rare taxa for which records were unavailable in some years in this range, as well as taxa that were not included in our phylogeny (see below). Our final dataset consists of 1,027,952 records from 49 species, which is the majority of odonate species (75 %) that have so far been recorded in Sweden (currently a total of 65; 43 dragonflies (Anisoptera) and 22 damselflies (Zygoptera)).

We obtained spatiotemporally matched estimates of spring mean surface temperature from the publicly accessible NOAA Global Forecast System Analysis (GFS ANL) database (https://www.ncdc.noaa.gov/data-access/model-data/model-datasets/global-forcast-system-gfs). The NOAA GFS ANL provides global weather data estimated four times per day gridded at .5° resolution (corresponding to approximately 25km in Sweden), using a system of satellites and globally-distributed weather stations. Comparison of these surface temperature estimates to direct temperature records from a weather station in Malmö, Sweden, obtained from the Swedish Meteorological and Hydrological Institute (https://www.smhi.se/en), indicates the GFS ANL data are an accurate representation of surface temperature (r > 0.99 between temperature records). For each record in our dataset of Swedish odonates, we obtained the mean surface temperature (GFS ANL database variable: tmp2m) from corresponding lat-long grid cell in the GFS ANL database for two time windows: the first 120 days of the year and the month of April. The first time window corresponds to a substantial period of pre-metamorphosis larval development and covers the last instar for all the species in our database, and so represents a long-term average temperature environment experienced during a key stage of development. The second time window represents a shorter time slice from late spring, just prior to the flight period of many early season species. We focused on mean temperature, because daily fluctuations in air temperature are unlikely to correlate strongly with water temperature, although average temperatures are a strong predictor of average water temperatures (Livingstone & Lotter 1998). We note that surface air temperatures are strong a strong predictor of water temperatures in freshwater bodies, particularly in the upper 1m of the water column (Livingstone & Lotter 1998) where odonate larvae are most abundant. Finally, we note that the scale and purpose of our analysis precludes use of a “sliding window” approach; Our interest is to compare developmental-temperature reaction norm variation across species (and so any such comparison requires use of similar or the same environmental cues), rather than to identify a specific time slice of spring temperature variation that is most important for a single species.

We obtained phylogenic information for all 49 species in our dataset, using a recently published large dated molecular phylogeny of the odonates (Waller & Svensson 2017). This tree is available publicly from the Odonate Phenotypic Database (http://www.odonatephenotypicdatabase.org/shiny/odonates/).

### Detailed Methods-Data Analysis

#### Long term field study of Ischnura elegans

We used a linear mixed effects model to assess phenological advance in the timing of the adult aerial period in *I. elegans* during our 18 year study period. This model featured the date (in ordinal day of the year; January 1 = day 1) of an observation of an individual as the response variable and year as a fixed effect. Our model also included a random intercept for population to account for spatial variation in phenology among the study populations; our model did not explore spatial autocorrelation, which we expect to be low as populations differ primarily in microenvironment that is uncorrelated with geography.

To infer selection on phenology, we seek a statistical description of individual fitness as a function of phenology, where the functional relationship between phenology and fitness varies with spring temperature cues. We modify the Lande-Arnold (Lande & Arnold 1983) equation to include temperature dependence in the form of directional selection on phenology:

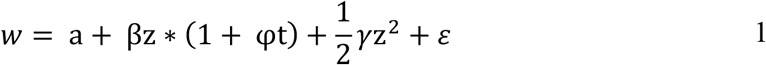

where 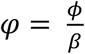, *ϕ* is an interaction term as estimable in a regression model, *w* is individual fitness, β is the linear selection gradient on phenology *z*, *t* is the temperature cue and *γ* is the nonlinear selection gradient on phenology. Setting the first derivative of equation 1 with respect to *z* to zero and solving yields the stationary point:

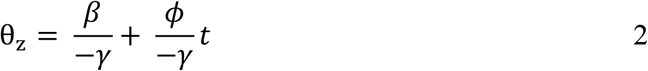

where θ_z_ is the optimum phenology (if selection is stabilizing, otherwise it is a fitness minimum). We note that this quadratic description of environmental-dependence of selection also approximates environmental dependence of a Gaussian fitness function (Chevin *et al.* 2015). The first part of the right hand side of equation 2 is an intercept equal to the fixed location of the optimum obtained from the standard Lande-Arnold equation (Phillips & Arnold 1989). The second term on the right hand side describes a temperature-dependent slope. Thus, the simple modification of the Lande-Arnold equation in 1 yields a statistical description of a linear relationship between temperature cue and optimal phenology, where the slope of the relationship between temperature and θ*_z_* is 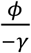, and represents a direct estimate of the temperature dependence of selection (*B*).

We fit equation 1 to our *I. elegans* field dataset using a linear mixed effects model with male copulation success as a response, i. e. a measure of sexual selection. For comparison, we also fit a simplified model, with directional selection only (i.e., no temperature-dependence or nonlinear terms). Although environmental effects on non-linear terms have not received theoretical or empirical attention (Chevin *et al.* 2015; Villemereuil *et al.* 2020), we nonetheless found no evidence for such effects and so focus on temperature effects on linear gradients. We focus on males only in this analysis, as copulation success is likely a more critical determinant of male fitness than it is for females; and also note that selection via male and female fitness components are strongly correlated across space and time in the wild (Gosden & Svensson 2008). We used absolute copulation success as a response, as this resulted in more consistent model convergence and eases interpretation; however, qualitatively equivalent conclusions were obtained using relativized (by subpopulation, season or globally) copulation success. We estimated β, *γ*, and *ϕ* from the fixed effect parameters (doubling the nonlinear regression coefficient; Stinchcombe *et al.* 2008), and included random (co)variation in β and *γ* among years to accommodate variation in selection independent of temperature and possible relationships between temperature and height of the fitness function emerging from equation 1. We fit an unstructured covariance matrix as we found no evidence for temporal autocorrelation in selection (ΔAIC = 135 for first order autoregressive covariance structure). We fit a gaussian-distributed model for consistency with theory and to provide estimates of the parameters in equation 1 (Chevin *et al.* 2015), although we also provide p-values from an equivalent mixed model assuming binomial error. Temperature was obtained as the mean of the first 120 days from the NOAA GFS ANL database for the corresponding grid cell of our study populations, with the exception of records from 2000-2005 for which we instead calculated this value from the Swedish Meteorological and Hydrological Institute data since the GFS data was unavailable for this period. We found close correspondence for the two years in which we had data from both of these sources (120 day mean, 2006: 1.061°C SMHI, 0.75°C NOAA; 2007: 5.89°C SMHI, 5.02°C NOAA).

#### Laboratory temperature experiment: raising damselfly families

We used a mixed effects model to estimate the relationship between thermal developmental environment of larvae (two temperature treatments) and emergence time from our laboratory experiment. Our model featured emergence day as the response variable, temperature treatment as a fixed effect, and random intercepts for tank and maternal full-sib family. We calculated phenotypic variances, additive genetic variances and heritability (*h^2^*), the latter as twice the family variance over the total variance of the random effects (the sum of tank, family, and residual variance components). We assessed statistical significance of *h^2^* using a likelihood ratio test of the family variance component. We also calculated standard error and 95% confidence intervals by empirically constructing the sampling distribution of *h^2^* by taking 1 million draws from a multivariate normal distribution centered on our original REML estimates of the variance components with covariance equal to the Hessian matrix of the fitted model (Houle & Meyer 2015). We note that our estimate of heritability is conditioned on the fixed effect of temperature treatment (Lynch & Walsh 1998).

#### Occurrence records of Swedish Odonata and phylogenetic comparative analyses

We used a bivariate mixed modelling approach (Phillimore *et al.* 2010; Phillimore *et al.* 2012) to infer interspecific variation in thermal reaction norms in phenology from our observational records of Swedish Odonata combined with paired estimates of temperature in the first 120 days of the year. The premise behind this approach is to use detrended covariation in temperature and phenology, within geographic grid cells, as an estimate of population mean reaction norms. From the same model, covariation in mean phenology and temperature across latitudinal geographic grid cells provides an estimate of the environmental dependence of selection (Hadfield 2016). We thus fit the bivariate model

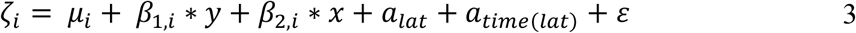

where *ζ*_i_ is the response vector of individual values of two variables: *i* = observed phenology or spring temperature. *β* is a fixed effect regression coefficient between continuous latitude (*y*) or year (*x*) and variable *i*. Two nested random effects are included in this model: *a_lat_* is a random effect describing (co)variation across binned latitude, and *a_time(lat)_* is a random effect describing co(variation) across binned annual seasons within latitudinal grids. For each *a* and the residual *ε* we estimated the full 2×2 unstructured covariance matrix 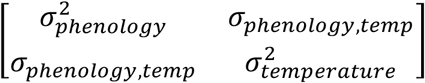 for a total of three 2×2 random effect covariance matrices. Thus, in this model space (latitude) and time (year) are included as both continuous fixed effects, as well as binned levels in nested random effects.

We focus on analysis of latitudinal spatial variation because Sweden spans a far greater latitudinal than longitudinal range, and corresponding species distributions within Sweden reflect substantial latitudinal span but little longitudinal span. Our dataset of odonate observations spanned latitudes from 55° – 69°, corresponding to the full latitudinal range of Sweden. We note, however, that longitudinal variation, when present, will be captured in variation in estimated spring mean temperatures for observations recorded from differing longitudes within the same latitudinal resolution, and thus is a source of residual within-bin variation accommodated in our model. For estimation of our mixed model random effects, we binned latitude by half degree increments, with the exception of extreme northern latitudes (over 62°) due to limited data from these sparsely populated Swedish regions. Our dataset contained 14 latitudinal bins with each species containing observations represented in at least 4 adjacent bins.

From this model we can interpret the ratio of the fixed effect parameter estimates, particularly 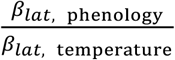 as approximating the environmental dependence of the optimum, *B* (Hadfield 2016). From the random effects, we can interpret *b* ∼ *σ*_phenology, temperature_/ 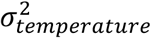 where *σ* and *σ*^2^ are the elements of the random effect covariance matrix estimated across time points within binned latitude (*a*_t*ime(lat)*_). Thus, *b* is the de-trended regression of phenology across temperature within latitudinal bins (Phillimore *et al.* 2010; Phillimore *et al.* 2012). This bivariate model was fit separately for each species in our dataset.

We then used the corresponding estimates of *B* and *b* in phylogenetic mixed models (Hadfield & Nakagawa 2010) to explore the relationship between species mean latitude, species mean phenology, and the strength and form of inferred phenological plasticity. We focus on species mean latitude because we were interested in how plasticity has evolved with Northern range expansion. These models included species as a random effect, with covariance among levels equal to the phylogenetic-variance-covariance matrix from our species level phylogeny, standardized to a depth of 1 (i.e., using the correlation matrix). These models also included a residual term. We accommodated uncertainty in our estimates of *B* and *b* in these phylogenetic mixed models by modelling the posterior variance of the species-specific estimates as a non-fitted random effect. We also estimated phylogenetic signal (“phylogenetic heritability”) in the inferred reaction norms *b* as the phylogenetic variance over the sum of the phylogenetic and residual variance (Pagel 1999; Hadfield & Nakagawa 2010) from a mixed model with *b* as the response, an intercept, and species as a random effect (in addition to the posterior variance in *b*, as above).

All bivariate and phylogenetic mixed models were fit by Bayesian MCMC using MCMCglmm in R and assuming weak priors. All other models were fit in SAS using the glimmix procedure. SAS and R scripts to reproduce our analyses and figures, as well as all raw data, are provided at: https://github.com/spdelisle/OdonatePhenology. All datasets have also been deposited as .csv files on Dryad https://datadryad.org/stash/share/FKHv4E_XmOX2f4yEi63pTibgzqmNFuXXhJLi_lnV3J0.

---

**Table S1.**
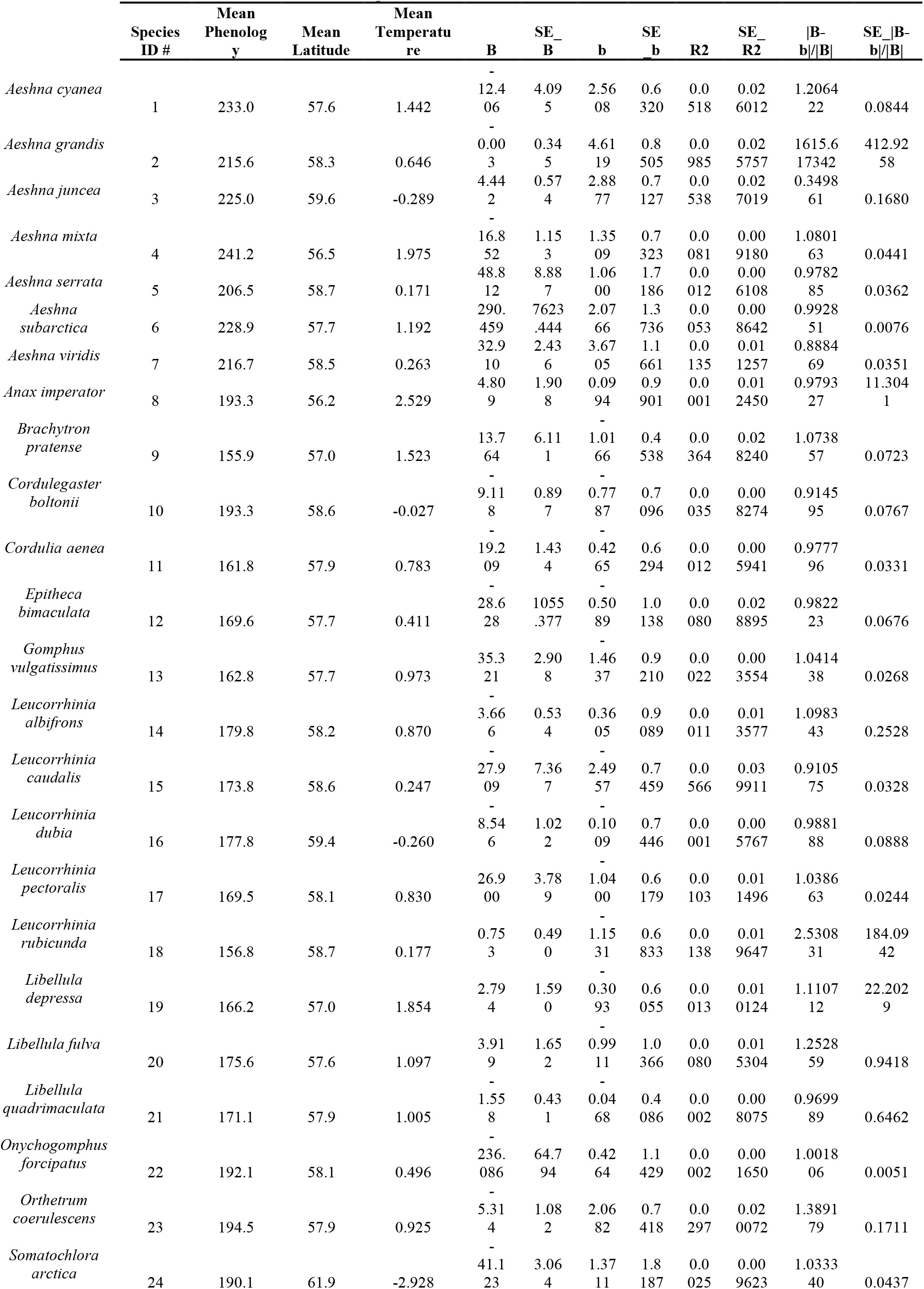
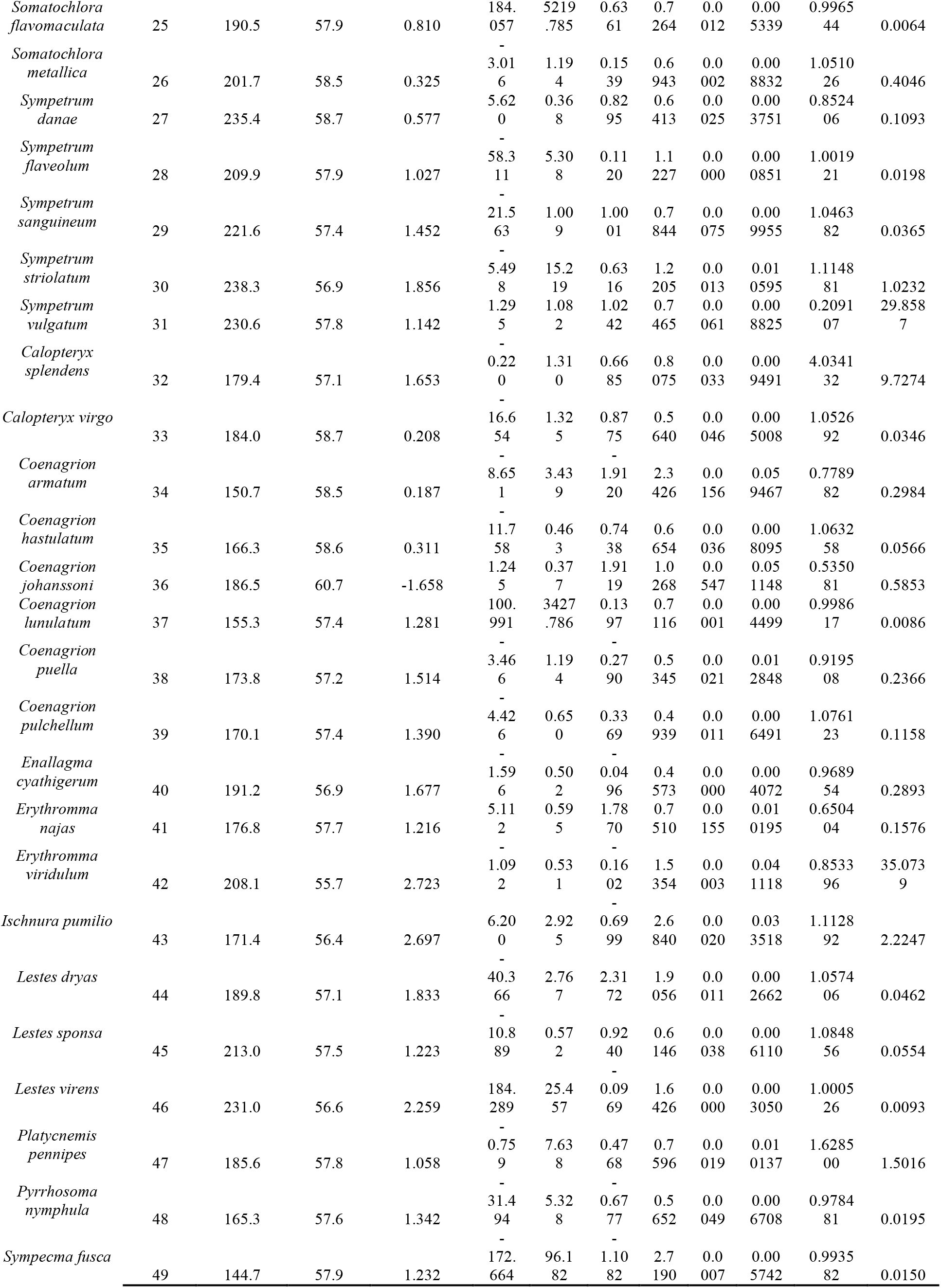
Summary statistics for 49 species of Swedish Odonata. B is the indirect estimate of the environmental dependence of the optimum, b is the indirect estimate of the temperature-phenology reaction norm. R2 is the variance in phenology explained by plasticity. Parameter values are posterior modes; standard errors are the square root of the

**Figure S1.**
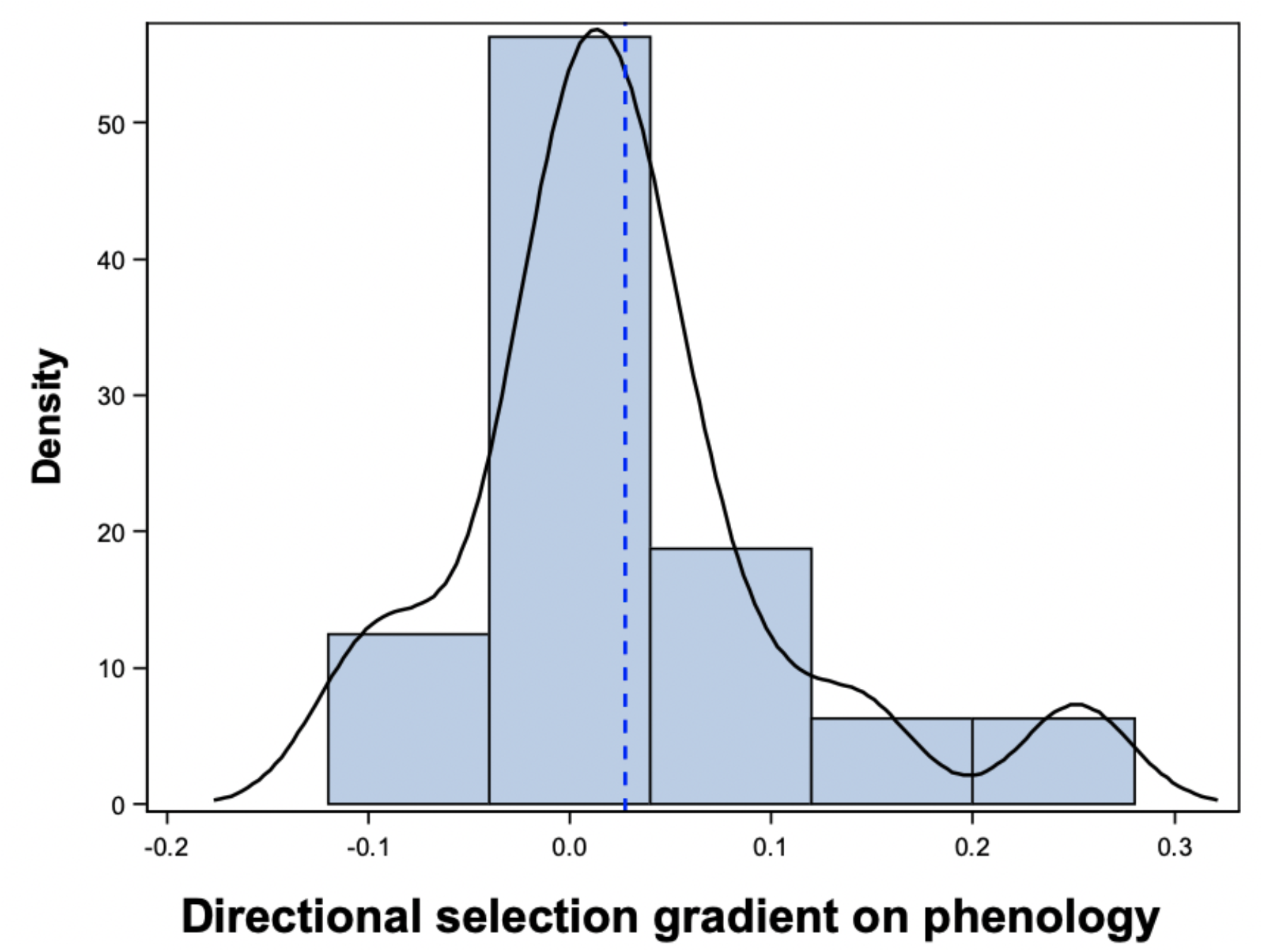
Variation in standardized directional selection gradients. Histogram shows BLUP estimates of annual directional selection on phenology, from a model with copulation success as response and globally-standardized (sd = 1) day as a fixed effect. Blue dashed line indicates the overall (main effect) estimate of directional selection across all 18 study years. Data come from a mixed effects model with random variation in slope and intercept among years.

**Figure S2.**
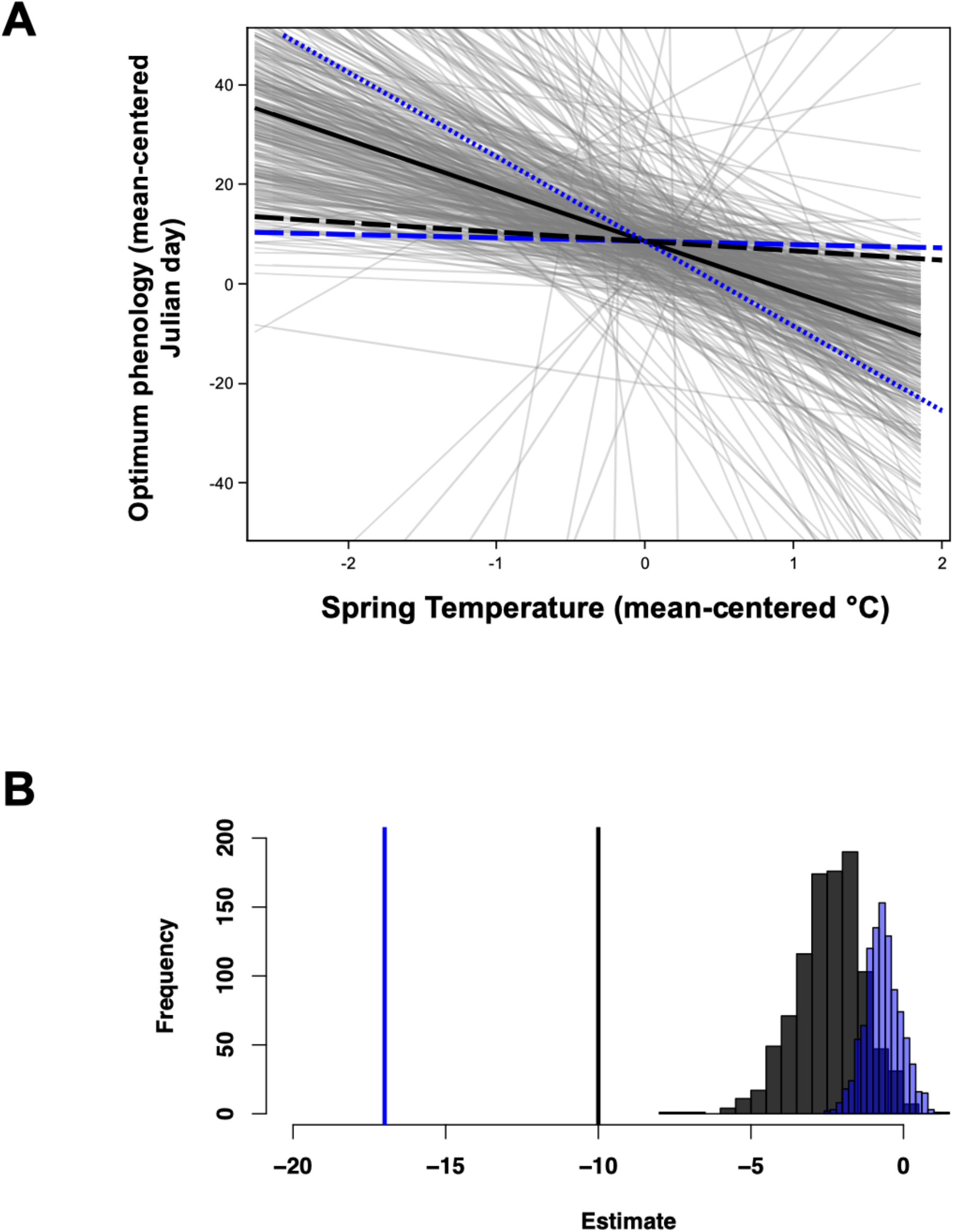
Comparison of parameter estimates from bivariate mixed effects reaction norm model and direct estimates of B and b. In panel A, Long-dashed blue and black lines are (respectively) the estimates of the population mean reaction norm, b, and the temperature-dependence of the optimum phenology, *B*, estimated from a bivariate mixed model with fixed and random effects of space and time (see text) using observational data from the Swedish Artportalen and NOAA GFS ANL. Other graph features are as in Figure 3C; although sampling effects result in slight differences in the grey lines showing error in B. Short-dashed blue line is the reaction norm b estimated in a laboratory experiment, solid black line is the estimate of B from individual fitness data from wild-caught damselflies. Panel B shows the posterior distributions of the observational estimates of *B* (black) and *b* (blue), along with the direct point estimates of these parameters. The direct estimate of *B* (*b*) differs significantly from the indirect measure of *B* (*b*).

**Figure S3.**
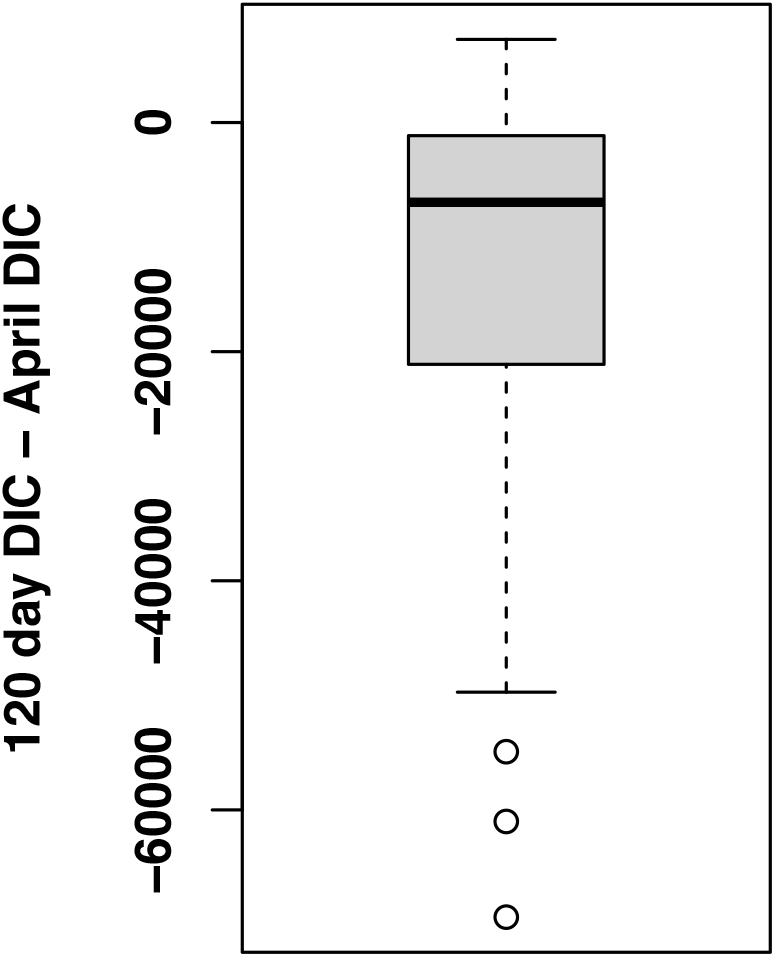
Comparison of model fits between identical bivariate mixed models. fit to temperature and phenology data using either the mean of the first 120 days of the year or the mean April temperature. The difference in Deviance shows that on average across the 49 species studied that the mean 120 days fit on average 14322 (95% CI: −19713.136 – −8932.425, P < 0.0001) DIC units better than mean April temperature. Note that these are identical datasets (equal N) and identical models, differing only in the values of mean temperature and so comparing the likelihoods in this way provides one approach to assessing fit.

